# Successful working memory linked to theta connectivity patterns in the hippocampal-entorhinal circuit

**DOI:** 10.1101/2022.09.08.507081

**Authors:** Jin Li, Dan Cao, Shan Yu, Haiyan Wang, Lukas Imbach, Lennart Stieglitz, Johannes Sarnthein, Tianzi Jiang

## Abstract

Working memory (WM) is the ability to actively maintain information for a short time and is central to human behavior. Rodent studies have proposed that hippocampal-entorhinal communication supports WM maintenance. However, the exact neural mechanisms of this interaction in WM remains unclear in humans. To address these questions, we combined machine learning analyses with intracranial electroencephalography (iEEG) recordings from the hippocampus and the entorhinal cortex (EC) in human participants, who maintained a set of letters in their WM. We found that WM maintenance was accompanied by elevated bidirectional hippocampal-EC information exchange via the theta band (2–8 Hz) and bidirectional cross-region theta-gamma phase-amplitude coupling (PAC). Further decoding analyses showed that the unidirectional inter-regional communication, with both theta oscillations in the hippocampus modulating EC gamma activity and theta band-coordinated information flow from the hippocampus, could decode correct performance at the level of participants. Taken together, our results demonstrate that theta functional coupling in the hippocampal-EC supports the maintenance of WM information via a specific pattern of frequency and direction. This connectivity-based coding could shed light on the neural mechanisms of WM processing.

**Significance:** Recent studies suggest a role for the hippocampus in working memory. How does the hippocampus coordinate with other brain regions to retain working memory information? The entorhinal cortex (EC) is the main gateway for information between the hippocampus and neocortex. To delineate whether (and how) the hippocampus and the entorhinal cortex interact during working memory and whether such interaction supports successful working memory, we used machine learning analyses of human intracranial EEG recordings while patients performed working memory tasks. Our results suggest that the human hippocampal-EC circuit supports working memory and is maintained in specific connectivity patterns, with a theta band (2–8 Hz)-coordinated unidirectional influence from the hippocampus to the EC. Our findings reveal that dynamic unidirectional interactions within the hippocampal-EC circuit underlie working memory and can contribute to a mechanistic circuit understanding of working memory.

## Introduction

Cognition critically depends on the ability to maintain information in an active and readily available state for a short time, which is typically ascribed to working memory (WM)(1). Studies found persistent single-neuron spiking (2-4) and elevated activity (5) in the hippocampus during WM maintenance. An increasing number of studies have pointed out that WM maintenance relies on multiple brain regions (6) and is supported by inter-regional communication (7, 8). Given this evidence for the role of hippocampus in WM and the distributed nature of WM, understanding the connectivity between the hippocampus and the rest of the brain could provide a crucial insight into the network involved in such a fundamental process. Then, one may ask how does the hippocampus interact with another/other brain area(s) during WM maintenance, and which brain area(s) contribute to this process?

The entorhinal cortex (EC) is a key candidate for such an area for the following reasons. First, studies consistently reported persistent spiking during WM maintenance of EC neurons in rats (9), monkeys (10) and human subjects (11). Second, the EC serves as an interface between the hippocampus and cortical and subcortical areas (12). Third, structural and functional hippocampal-EC interactions have been widely reported as follows: At the anatomical level, tracing studies have uncovered structural connections between the two brain areas (13, 14). At the functional level, the hippocampus receives multiple sensory information from the EC and feeds memory-associated activity back to the EC (15, 16). At the behavioral level, rodent studies have indicated that hippocampal-EC communication supports WM maintenance. For instance, electrophysiological recordings have observed synchronized oscillations during correct, but not incorrect, WM execution in rats (17). In addition, inhibition of the hippocampal-EC circuit using optogenetic techniques (17) or a transgenic model (18) in rodents resulted in significant WM impairments. However, these animal studies, which suggest that the hippocampus-EC circuit supports WM, have not been validated in humans. Investigating the dynamic oscillation of circuit communication in humans has been hampered by limitations such as poor spatial and temporal resolution, which are associated with using noninvasive methods.

If the hippocampal-EC circuit contributes to WM, a reasonable next question would be, how does the oscillatory communication between the hippocampus and the EC mediate WM? Specifically, is there a certain oscillatory mode in a specific frequency or direction that underlies successful WM? Theta and gamma oscillations, representing continuous and transient oscillatory phenomena (19, 20) respectively, are the main types of rhythmic field activity generated in the hippocampus and EC in both rats and humans (21-23). Studies suggested that synchronous theta oscillations and theta-gamma phase amplitude coupling rhythms facilitate inter-regional information communication and are implicated as neural correlates of WM (24, 25). Therefore, we hypothesized that theta oscillations mediate functional connectivity between the hippocampus and the EC by information flow via theta oscillations and by coupling inter-regional theta-gamma oscillations.

Leveraging the high spatiotemporal resolution of iEEG recordings and the analytical power of multivariate machine-learning analysis, we tested the hypothesis that theta and gamma oscillations cooperatively facilitate hippocampal-EC interactions to support the maintenance of WM information in humans. We recorded iEEG data simultaneously from the hippocampus and the EC in 13 epilepsy patients while they performed a modified Sternberg task (4, 5). Our goal was to address the following questions: (a) How do hippocampus and the EC interact while humans perform a WM task? (b) Do these interactions support successful WM maintenance? And (c) which specific oscillatory modes of interregional communication, including frequency and directionality, support successful WM maintenance?

## Results

### Task, behavior and recording channels

Thirteen patients with drug resistant epilepsy (6 female) performed a modified Sternberg WM task during an invasive presurgical evaluation. In this task, the items were presented simultaneously rather than sequentially, thus separating the encoding period from the maintenance period. In each trial, the participant was asked to memorize a set of letters presented for 2 s (encoding). After a delay (maintenance) period of 3 s, a probe letter was presented and the participant responded whether the probe letter was identical to one of the letters held in memory (retrieval) (**Fig. 1(A)**). The average accuracy was 92.0% ± 3.3% (range 86.1%-97.6%). The mean response time for correct trials was 1.44 ± 0.38 seconds. Hence, the participants performed well in the task. Local field potentials were recorded simultaneously from depth electrodes implanted in the hippocampus and the EC (**Fig. 1(B)**). In total from all participants, 87 channels in the hippocampus and 46 channels in the EC were included in the subsequent analysis (see the details in **Methods**).

**Fig. 1.**
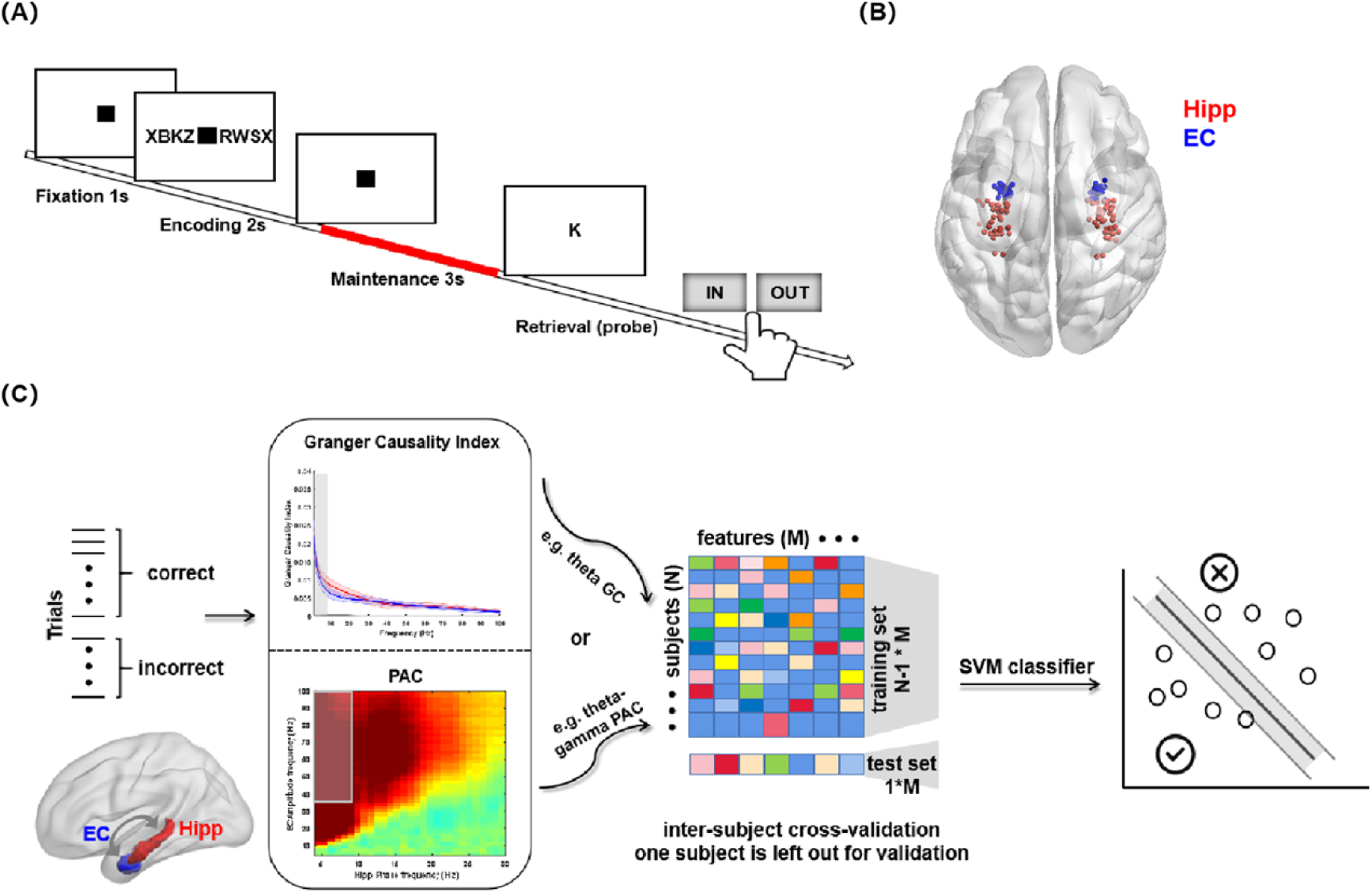
Working memory task, recording sites, and schematic of multivariate pattern analysis (MVPA) using patterns of information flow and cross-frequency coupling between the hippocampus and the entorhinal cortex. (A) An example trial of the task. Each trial consisted of a set of consonants (encoding 2 s), followed by a delay (maintenance 3 s). After the delay, a probe letter was shown, and the subjects indicated whether the probe was or was not shown during the encoding period (retrieval 2 s). (B) Channel location across subjects in MNI152 space (see Methods). Recording regions of the hippocampus (Hipp) and entorhinal cortex (EC) are indicated by different colors (red, Hipp; blue, EC). (C) Schematic of the multivariate pattern analysis. Granger causality index (GC) as well as phase-amplitude coupling (PAC) was estimated between the Hipp (red) and the EC (blue) across the correct and incorrect trials, separately. The patterns of the GC (e.g., theta GC) and PAC (e.g., theta-gamma PAC) were separately used to train the support vector machine (SVM) classifier to classify the WM performance (correct or incorrect). Specifically, we converted the features to a feature vector (M), combined all the subjects’ (N) data, fed them into a linear SVM classifier, trained the classifier on the data from N-1 subjects (N-1*M), and tested it on the remaining subject (1*M). We used inter-subject cross validation by leaving one subject out for validation and replicated the classification. The accuracy was used as the performance metric. For further details, see the Methods.

### Directional information transfer from the hippocampus to the EC at the theta band decoded the WM outcome

First, we used a frequency-domain Granger causality (GC) analysis to quantify the inter-regional directional influence, which measures the degree to which the signal from a region can be better predicted by incorporating information from another signal in a specific frequency band (26). We computed the spectral GC index from the hippocampus to the EC, as well as in the reverse direction, during maintenance for trials with correct and incorrect performance, separately. As presented in **Fig. 2(A)**, the GC index for both directions (red, from hippocampus to EC; blue, from EC to hippocampus) was above the threshold for significance (grey lines denote the thresholds), in both correct and incorrect trials. No difference in the GC index across 1-100 Hz was found between the two directions (cluster-based permutation test, *p* > .05). This finding revealed bidirectional information exchange across a broad frequency band within the hippocampus-EC circuit during WM maintenance.

**Fig. 2.**
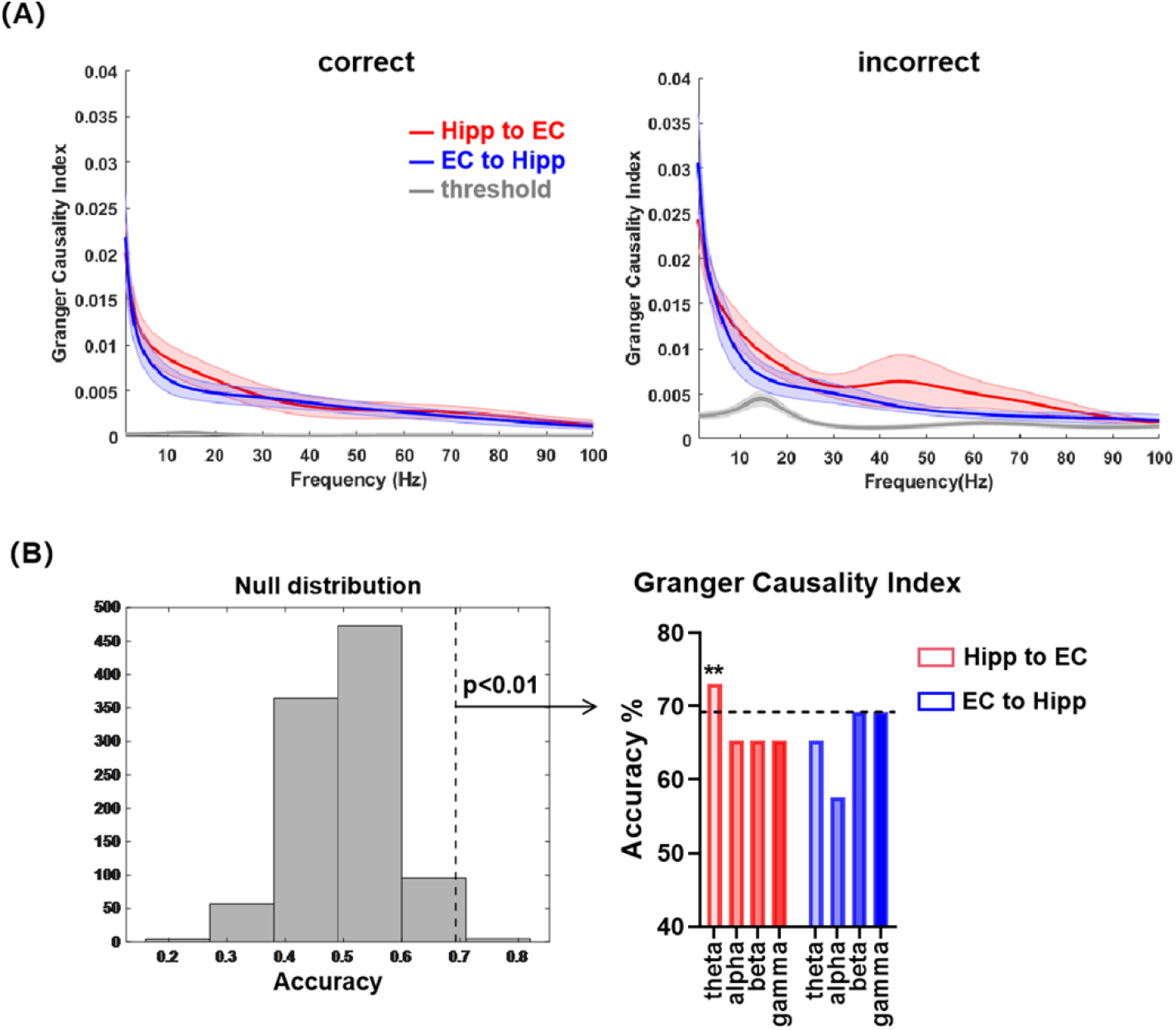
Information flow between the hippocampus and entorhinal cortex, and decoding accuracy in the GC-based decoding analyses during WM maintenance. (A) Average GC value between the Hipp and the EC (red, Hipp to EC; blue, EC to Hipp) across all subjects during maintenance for the correct (left) and incorrect (right) trials, separately. No difference in GC was found between the two directions (cluster-based permutation test, *p* >.05) across 1-100 Hz. Shaded areas, SEM. (B) Decoding accuracy based on the GC values from two directions (red, Hipp to EC; blue, EC to Hipp) in the theta (2-8 Hz), alpha (8-13 Hz), beta (13-30 Hz), and gamma (30-100 Hz) frequency bands during WM maintenance. The left panel shows the null distributions of the maximum statistics for all the decoding analyses, which were created using classifiers with randomized training labels. The threshold of significance is marked with a vertical dashed line (69.23%, *p* < .01). The right panel shows the decoding accuracy values for each direction and frequency. Those above the threshold (horizontal dashed line) were deemed statistically significant. Only the decoding accuracy by the theta band features from the Hipp to EC significantly decoded the WM performance, which is labeled with asterisks (**, *p* < .01).

Next, we addressed the question of whether inter-regional causal interactions supported successful WM. Support vector machine (SVM) classifiers show good generalization performance for high dimensional data and have been used widely for classifying scalp EEGs (27) and have recently been successfully utilized for classifying magnetoencephalography (8) signals. Hence, we used a linear SVM classifier here to decode WM performance on the subject level (correct or incorrect), with the extracted GC values from two directions as features (as shown in **Fig.1(C)**). The data from all the trials in all the participants (N=13) were used in the classification, which was cross-validated by leaving one participant out for validation. This means that we trained the model with data from N-1 participants and tested the model using the data from the one left out participant. The goal of the whole classification process was to classify the WM outcome. The statistical significance of the classification accuracy was determined by comparing the original accuracy with a null distribution created by using a randomized classifier by permuting the labels 1000 times (for further details of the statistical analyses, see **Methods**).

The extent to which different frequencies underlie the hippocampal-EC interaction in WM remains an open question because the connectivity has not been fully mapped across frequency bands, although theta communications within this circuit are widely reported to support cognitive function (28). Therefore, we separately extracted the GC values at the theta (2-8 Hz), alpha (8-13 Hz), beta (13-30 Hz), and gamma (30-100 Hz) bands from two directions: from the hippocampus to the EC and the reverse direction for the correct and incorrect trials. We then converted the GC features at a specific frequency band to a feature vector and combined the data from all the participants to decode the WM outcome using the SVM classifier (for details, see **Methods**). The results indicated that only the decoding accuracy using features in the theta band from the hippocampus to the EC exceeded the threshold of significance (permutation test, *p* < .01, see **Fig. 2(B) left**). The decoding accuracies of the other bands (alpha, beta, and gamma) from the hippocampus to the EC as well as the features within all frequency bands from the EC to the hippocampus were not above the threshold for significance (permutation test, all *p* > .01). These results suggest that successful WM maintenance is associated with directional information transfer from the hippocampus to the EC in the theta band in individual participants.

### Distinct theta phase encoded information from the hippocampus and EC

Phase-amplitude coupling (PAC) serves as an important mechanism for coordinating interregional information transfer because high frequency amplitudes are modulated by low frequency phases (29). It has been suggested that PAC, especially theta-gamma PAC within the hippocampus, reflects a neural correlate for WM maintenance in humans (5, 30). We computed the cross-regional PAC in both phase-amplitude combinations between the hippocampus and the EC for the correct and incorrect trials, separately. The interregional influence was quantified as the coupling of low-frequency (2-30 Hz) phases from one region with high frequency amplitudes (30-100 Hz) to another. The raw PAC was *z-*scored against surrogate distributions via bootstrapping (see **Methods**). Based on the GC results, we focused on the theta (2-8 Hz)-gamma (30-100 Hz) PAC between the hippocampus and the EC (gray box in **Fig. 3(A)**). For the incorrect trials, both directions had modulation effects between the hippocampus and the EC; these effects peaked at 30 Hz (**Fig. 3(A) right**). In addition, both directions of the *z*-scored PAC were significantly higher than 0 for both the correct and incorrect trials (cluster-based permutation test, *p* < .05), a finding which indicated real inter-regional coupling effects.

**Fig. 3.**
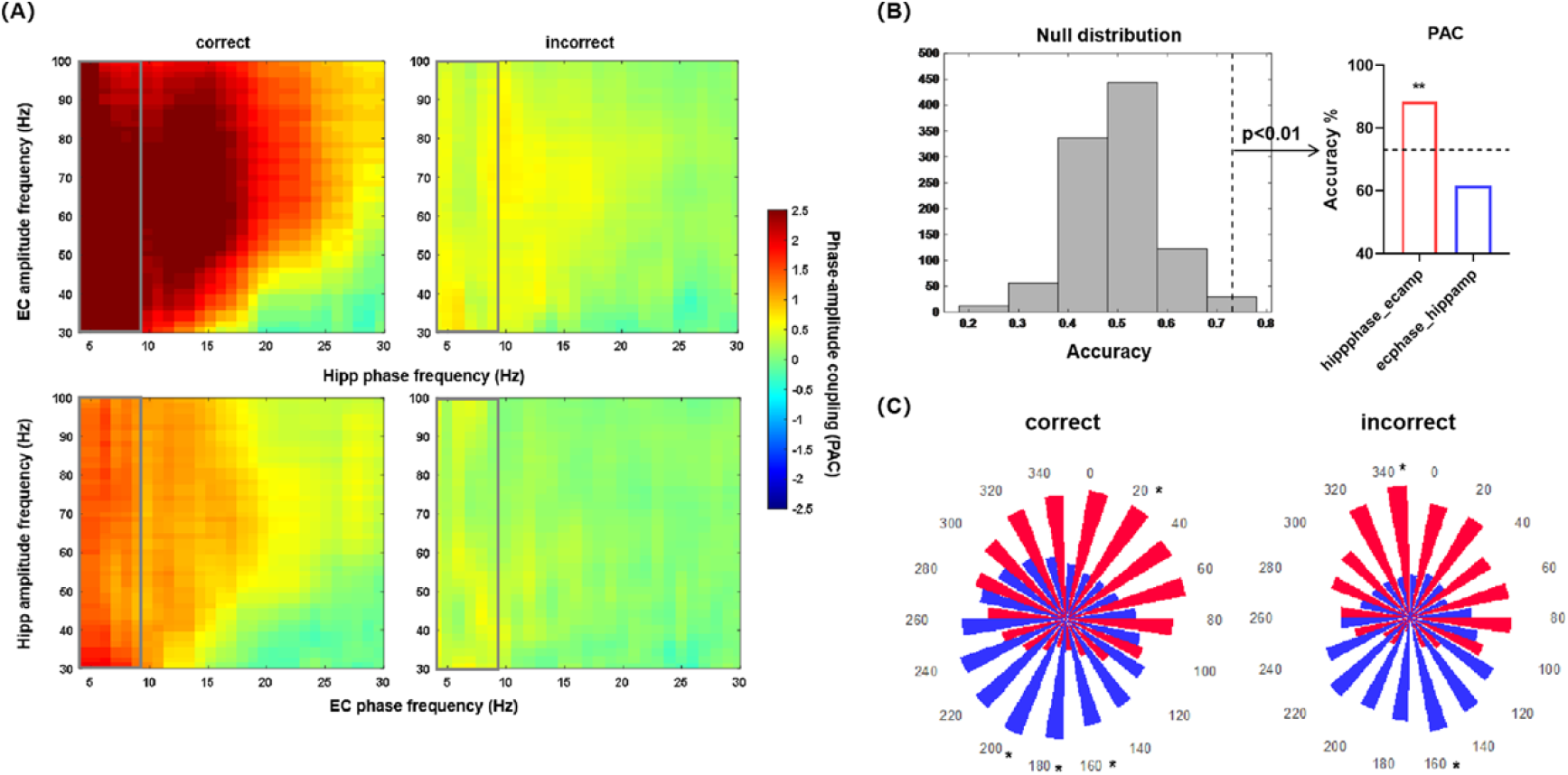
Cross-frequency coupling between the hippocampus and entorhinal cortex, and decoding accuracy in the coupling-based decoding analyses during WM maintenance. (A) Average cross-frequency PAC (*z*-score) between the Hipp and the EC across all subjects for the correct and incorrect trials, separately. The top row depicts the Hipp phase modulating the EC amplitude and the bottom row depicts the EC phase modulating the Hipp amplitude. Gray boxes delineate theta phase (2-8 Hz)-gamma amplitude (30-100 Hz) coupling. Both directions had a significant inter-regional coupling effect during maintenance (cluster-based permutation test, *p* < .05) in both correct and incorrect trials. (B) Decoding accuracy based on the theta-gamma PAC values from two directions (red, Hipp modulating EC; blue, EC modulating Hipp) during WM maintenance. The left panel shows the null distributions of the maximum statistics for the two decoding analyses, which were created using classifiers with randomized training labels. The threshold of significance is marked with a vertical dashed line (99% of null distribution, 73.08%). The right panel shows the decoding accuracy values for each direction. Those above the threshold (horizontal dashed line) were deemed statistically significant. Only the decoding accuracy using the PAC features from the Hipp modulating the EC significantly decoded the WM performance, which is labeled with asterisks (**, *p* < .01). (C) Computation of the theta-gamma interactions for the correct and incorrect trials. Phase-amplitude distributions were constructed for both directions (red, EC gamma amplitudes distributed across the theta phase bins of Hipp, Hipp-EC; blue, Hipp gamma amplitudes distributed across the theta phase bins of EC, EC-Hipp). For both the correct (left) and the incorrect trials (right), the gamma activity from both the Hipp (red) and the EC (blue) was modulated by the theta oscillations around the trough (Rayleigh test, for the correct trials, *p*_*EC*_ = 4.38 × 10^−13^, *p*_*Hipp*_ = 4.39 × 10^−13^; for the incorrect trials, *p*_*EC*_ = 4.39 × 10^−13^; *p*_*Hipp*_ = 4.39 × 10^−13^). Significant differences between the two directions were found at several phase bins (paired *t* tests, all *p*s < .05) for both correct and incorrect trials. * denotes *p* < .05.

The theta phase-gamma amplitude *z*-scored PAC values were extracted from two directions: the hippocampal phase modulating the EC amplitude and the reverse direction for the correct and incorrect trials, separately. Again, these *z*-scored PAC features were fed into the linear SVM classifier to decode the WM performance. The procedure was similar to that in the GC section except that the features were replaced with PAC values. Only the features from the theta phase of the hippocampus that entrained the gamma amplitude of the EC could significantly decode the WM performance (permutation test, *p* < .01, see **Fig. 3(B)**), while the decoding accuracy from the reverse direction did not reach the threshold (permutation test, *p* > .01). These results indicate that the hippocampus and the EC use a direction-specific PAC to optimize WM maintenance.

In addition, given that low-frequency oscillations are thought to organize cell assemblies to maintain WM information (31), we also tested whether a specific phase of the theta oscillation was co-modulated with the gamma amplitude in a behaviorally relevant manner. The distributions of the gamma amplitudes across 18 phase bins were calculated for both directions. As shown in **Fig. 3(C)**, the gamma activity from both the EC (red) and the hippocampus (blue) was modulated by the theta oscillation around the trough for the correct trials (left, Rayleigh test, *p*_*EC*_ = 4.38 × 10^−13^; *p*_*Hippocampus*_ = 4.39 × 10^−13^) as well as for the incorrect trials (right, Rayleigh test, *p*_*EC*_ = 4.39 × 10^−13^; *p*_*Hippocampus*_ = 4.39 × 10^−13^). Next, we compared the distributions of the gamma activity between the hippocampus and the EC at each phase bin using paired *t* tests and applied false discovery rate (FDR) approach for multiple comparisons. In summary, these results not only confirmed the directional interactions observed from the Granger causality analysis but also suggested that such a directional influence of the theta oscillation could shape the gamma amplitudes. High gamma oscillations are typically phase locked to the trough of the theta oscillation. These findings suggest that a distinct phase of the theta oscillation was modulated by the gamma activity in a directionally relevant manner and that this underlies WM maintenance.

### Local neural oscillations from the hippocampus and EC did not support successful WM maintenance

The above analysis revealed that the WM outcome can be decoded by the hippocampal-EC interactions in the theta band and in the theta-gamma couplings. Then, we asked whether local activity in the hippocampus and the EC contribute to successful WM maintenance. To address this question, we calculated the time-frequency power for each channel within the hippocampus and EC for the correct and incorrect trials, separately. The power outputs were *z*-scored against the pretrial baseline distributions to assess the significance of the task-induced power effects per trial (see **Methods**). We extracted the *z*-scored power within the hippocampus and EC at the theta (2-8 Hz) and gamma (30-100 Hz) bands across the maintenance period for each participant, and compared them between the correct and incorrect trials using paired *t* tests. No difference in *z*-scored power was found for any frequency band within each region (paired *t* tests, all *p* > .05, **Fig. 4(A)**). Next, the *z*-scored power values for the theta and gamma bands within the hippocampus and EC were used as features in the SVM classifier. No significant results (**Fig. 4(B) right**) were found for any of the bands and regions; that is, none exceeded the threshold of significance (69.23%, 99% of the null distribution, **Fig. 4(B) left**). This implies that local neural activity from the hippocampus and EC did not support the differentiation of the WM performance.

**Fig. 4.**
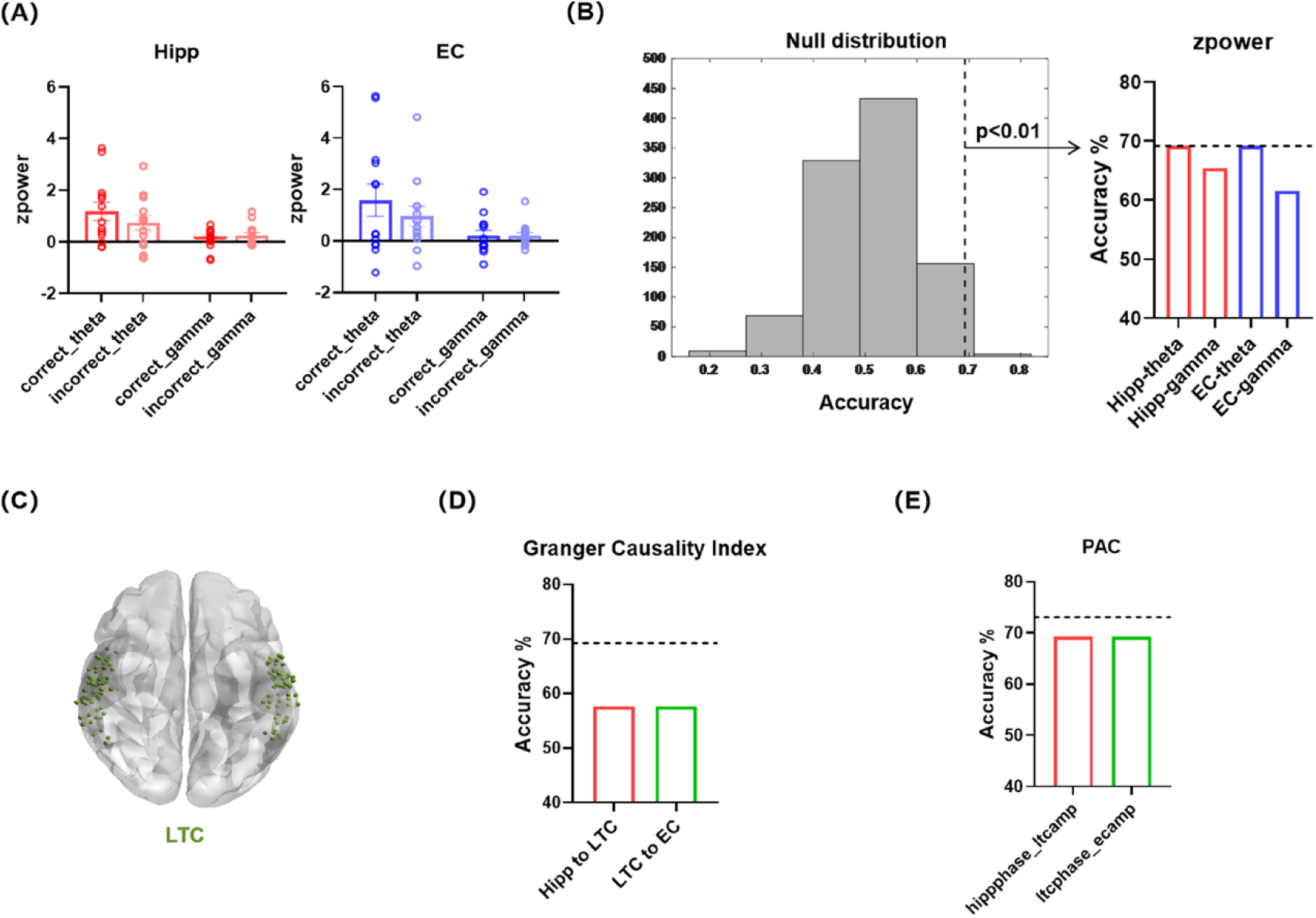
Control analyses based on the spectral power of the Hipp and EC, and decoding analyses with GC and PAC of the lateral temporal cortex (LTC) as a control site. (A) Average *z*-scored power in the theta (2-8 Hz) and gamma (30-100 Hz) bands during maintenance for the correct and incorrect trials within the Hipp (left) and the EC (right). No difference was found between the correct and the incorrect trials (paired *t* test, all *p*s> .05) in each region and frequency band. Dots denote individual subjects. (B) Decoding accuracy based on the *z*-scored power values of the Hipp (red) and EC (blue) at theta (2-8 Hz) and gamma (30-100 Hz) frequency bands during WM maintenance. The left panel shows the null distributions of the maximum statistics for all decoding analyses, which were created using classifiers with randomized training labels. The threshold of significance is marked with a vertical dashed line (99% of null distribution, *p* < .01). The right panel shows the decoding accuracy values for each region and frequency. Those above the threshold (horizontal dashed line) were deemed statistically significant. The decoding accuracy based on the *z*-scored power did not exceed the threshold (69.23%). (C) Recording sites in the LTC, which are indicated by all contacts across subjects (green), were used as controls. (D) Decoding accuracy based on the GC values from the Hipp to the LTC (red) and the LTC to the EC (green) in the theta (2-8 Hz) frequency band during WM maintenance. The decoding accuracy values for each direction did not exceed the threshold (69.23%) used in the decoding analyses with the GC features from the Hipp and EC. (E) Similarly, decoding accuracy based on the theta phase-gamma amplitude PAC values from the Hipp phase -LTC amplitude (red) and the LTC phase - EC amplitude (green) during WM maintenance. The decoding accuracy values for each direction were not statistically significant.

### Decoding analyses with the lateral temporal cortex as a control site

Furthermore, to confirm that the contribution of hippocampal-EC interaction to WM maintenance was not a general effect, we chose the lateral temporal cortex (LTC, all recording sites presented in **Fig. 4(C)**) as an anatomic control region. We repeated the above SVM analysis within the hippocampal-EC circuit, each time substituting the feature of one region for that in the LTC. Our results found that none of the estimated decoding accuracies was above the threshold of significance, regardless of whether we used the theta GC (**Fig. 4(D)**) or the theta - gamma PAC feature patterns (**Fig. 4(E)**) between the hippocampus/EC and the LTC.

## Discussion

In this study, we found that interregional communication between the hippocampus and the EC uses specific oscillatory modes of frequency and direction to support successful WM maintenance. In particular, significant connectivity-based decoding results were found in hippocampal driven information transfer via the theta band and via PAC between the theta phase of the hippocampus entraining the gamma amplitude of the EC. These findings provided direct neural evidence of hippocampal-EC interactions during WM maintenance in humans and links a specific inter-regional pattern to successful WM performance.

These interregional oscillatory dynamics are consistent with known structural and functional connections between the hippocampus and the EC. Anatomical studies found that the EC sends projections to and receives monosynaptic input from the hippocampus (13, 32). Optogenetic inhibition of this circuit in mice resulted reduction in both inter-regional connectivity and the correct execution of WM-guided behavior (17). Our results are thus consistent with animal literature suggesting the contribution of hippocampal-EC communications to WM processing and extended these findings to humans. To date, only a handful of human studies have collected direct intracranial data on both the hippocampus and the EC during WM processing, and all looked at each region separately rather than at their connectivity (2, 3). We found that the WM outcome could only be decoded by the connectivity, not by the local activity within the hippocampal-EC circuit, suggesting that the hippocampus and the EC orchestrate to better store WM information. This underscores the necessity of considering connections rather than local activities in future studies to understand the neural mechanisms of WM.

We found that unidirectional influences from the hippocampus to the EC contribute to successful WM. Several studies indicated that the hippocampus receives sensory information from the EC. Then, the hippocampus processed the information and returned memory information to the EC (15, 16). During WM maintenance, the stimulus was not visible on the screen and needed to be held in memory. Therefore, memory information and not sensory information was needed. Thus, it is reasonable that the hippocampal outflow plays a key role in successful WM. Future neural modulational studies aimed at enhancing the WM performance should consider a more precise model of the specific direction within the hippocampal-EC circuit. In addition, the hippocampal outflow during WM maintenance might also accompany the memory transfer from short-term storage in the hippocampus to long-term storage in the neocortex, that is, in the process of memory consolidation (33). Recent studies found that memory consolidation may start as early as at the end of encoding (34, 35). In agreement with this, hippocampal outflow during the post-encoding period could decode subsequent memory performance (34), and WM maintenance contributes to long-term memory performance (36). Taken together, our results may have implications for understanding long-term memory consolidation.

The SVM analysis showed that theta band-driven information flow is instrumental in maintaining WM information and in achieving correct performance. Theta oscillatory plays a crucial role in enabling inter-regional interactions (37) and information processing during WM (24, 38, 39). Computational models have suggested that theta oscillation coordinates the proper timing of interactions between the hippocampus and the EC (40). Accumulating evidence indicates that the communications within the hippocampal-entorhinal circuit via the theta band constitute a common substrate for episodic memory (28) by coordinating neural ensembles and facilitating synaptic plasticity (41-43). We inferred that these neural mechanisms not only support episodic memory but also facilitate information processing during WM maintenance.

In subsequent analysis with cross-frequency coupling features, WM outcome could be decoded by theta-gamma PACs. Theta and gamma are two major oscillations of local field potentials in both the hippocampus and the EC during a WM task in rodents and humans (44, 45). Theta-gamma PACs have been suggested as modulating synaptic plasticity (46) and flexibly organizing complex mnemonic information (47). An influential theory model proposed that several WM items are activated in different gamma cycles during every cycle of the theta oscillation (48), and enhanced theta-gamma coupling was observed for correctly identified WM items (49). Recent human studies using iEEG recordings observed that the PAC, with theta oscillations in the hippocampus modulating the EC’s gamma activity, supports episodic memory (50). Our study extends this finding to WM processing.

In summary, our results provide direct evidence that successful WM maintenance is supported by the unidirectional influence from the hippocampus to the EC via the theta band. We have extended previous knowledge of the contribution of the hippocampal-EC circuit on WM in animals to humans. Such work will reveal the basic mechanisms of WM while also advancing translational interventions to treat human with WM disorders.

## Materials and methods

### Participants

We utilized data from 13 adult human participants (mean ± SD [range]: 35±13 [18-56]; 6 females) in this study. All participants were implanted with depth electrodes (1.3 mm diameter, 8 contacts of 1.6 mm length, and 5 mm spacing between contact centers; Ad-Tech, Racine, WI, www.adtechmedical.com) in the medial temporal lobe for evaluation of the surgical treatment of epilepsy. Electrode placement was exclusively guided by clinical needs. There were no seizures recorded during any of the epochs, and any epochs with interictal epileptiform activity were excluded from analysis.

### Ethics statement

All the participants provided written informed consent before testing. This study had been approved by the relevant institutional ethics review board (Kantonale Ethikkommission Zürich, PB 2016-02055), and was also in agreement with the Declaration of Helsinki.

### Experimental description

WM was tested using a modified Sternberg task in which the encoding of memory contents, maintenance, and recall were temporally separated (**Fig. 1(A)**). Each trial started with a fixation period (1 s) followed by the stimulus for 2 s. The memory content consisted of a set of letters at the center of the screen. After the disappearance of the stimulus, the maintenance interval was followed by a fixation square (3 s). Then, the participants responded with a button press (“IN” or “OUT”) to indicate as quickly as possible whether the probe was part of the stimulus when a probe was presented. After the response, the participants were encouraged to relax, and the next trial was initiated. The participants performed 50 trials per session, which lasted approximately 10 min. The order of the stimuli was fully random across sessions. During the recording period of several days, several participants were asked to perform more than one session of the task up to a total of seven sessions.

### Channel localization and selection

The channels were localized using postimplantation computed tomography (CT) scans and postimplantation structural T1-weighted MRI scans. For each patient, the CT scan was co-registered to the postimplantation scan, as implemented in FieldTrip toolbox (51). The channels were visually marked on the coregistered CT-MR images. Channel positions were verified by the neurosurgeon (L.S.) after merging pre-operative MRI with postimplantation CT images of each individual patient in the plane along the electrode (iPlan Stereotaxy 3.0, Brainlab, München, Germany). Channel locations in native space for each patient were projected to MNI space.

Each participant’s hemisphere had a maximum of three electrodes targeting the anterior and posterior hippocampus and the entorhinal cortex. Targeted regions and hemispheres varied across participants for clinical reasons and included the hippocampus in the left (*n* = 12) and right (*n* = 13) hemispheres and the entorhinal cortex in the left (*n* = 12) and right (*n* = 11) hemispheres. We selected the two most medial channels on each electrode targeting the hippocampus or the entorhinal cortex, as was done in previous studies (52, 53). This procedure was used to minimize inter-individual variability, which would be higher if different numbers of channels had been selected across participants. The final number of selected channels in each region for each participant is listed in **Table 1**. We included only ipsilateral channel pairs in the analysis. The final dataset contained 87 channels in the hippocampus and 46 channels in the entorhinal cortex across all patients. There were 6.7±1.5 (range 4-8) channels per patient in the hippocampus and 3.5±0.9 (range 2-4) channels per participant in the entorhinal cortex. We presented all channels from the hippocampus and the entorhinal cortex across all participants in **Fig.1(B)** and plotted them using BrainNet Viewer toolbox (54) in MATLAB (MathWorks, Natick, MA).

**Table 1.**
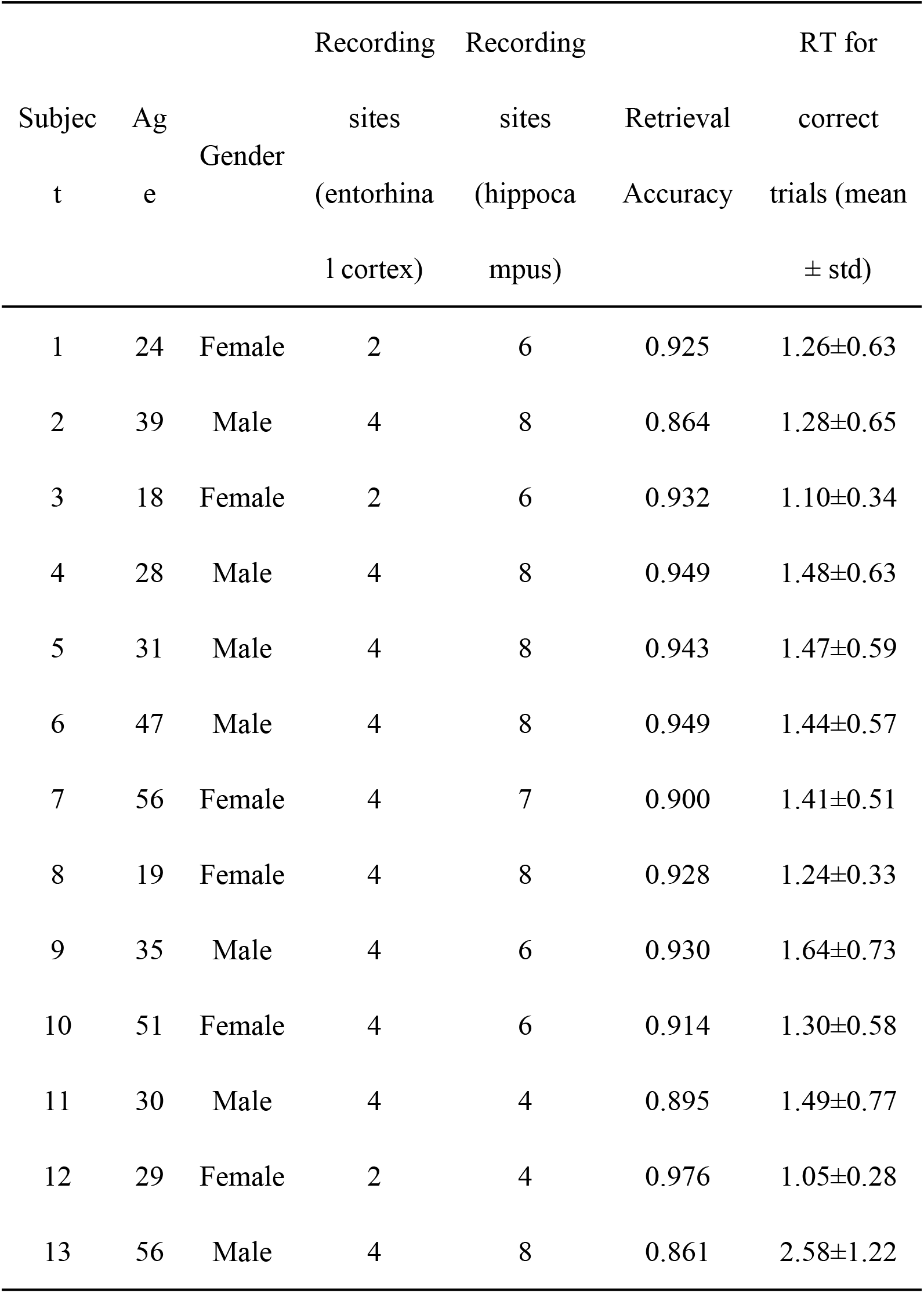
Subject characteristics.

In addition, we chose the lateral temporal cortex as an anatomic control site. As a previous study did (55), in each participant, the most two central channels in the gray matter in the lateral temporal cortex were on the same lead as the hippocampus/entorhinal cortex electrodes.

### Data acquisition and preprocessing

Intracranial data were acquired using a Neuralynx ATLAS recording system, sampled at 4 kHz, and analog-filtered above 0.5 Hz against a common intracranial reference. After data acquisition, neural recordings were down sampled to 1 kHz and band-pass filtered from 1 to 200 Hz using the zero-phase delay finite impulse response (FIR) filter with a Hamming window. Line noise harmonics were removed using a discrete Fourier transform. The filtered data were manually inspected to mark any contacts or epochs containing epileptiform activity or artifacts for exclusion. The data were then re-referenced to a bipolar reference. The continuous data were segmented into 8 s trials with a 1 s fixation period as the baseline, 2 s encoding period, 3 s maintenance period, and 2 s retrieval period. We focused on the maintenance period. The trials with residual artifacts were rejected after visual inspection. The participants’ responses defined the correct and incorrect trials. Preprocessing routines were performed using the FieldTrip toolbox (Oostenveld et al., 2011) and customized scripts for MATLAB (MathWorks, Natick, MA).

### Granger causality analysis

We examined the information directionality of the hippocampus-EC synchronization using spectral GC, which quantifies the prediction error of the signal in the frequency domain by introducing another time series. For each channel pair, the trial-wise mean was subtracted from each trial before being fit to an autoregressive model and computing the spectral GC. We then applied the Multivariate Granger Causality Matlab Toolbox (56) based on the Akaike information criterion to define the model order for each pair. The GC index was computed across 1-100 Hz (in steps of 0.25 Hz) for both directions (from the hippocampus to the EC and the reverse direction) during maintenance for the correct and incorrect trials separately. Then we created a null distribution by randomly swapping the signal between the channels 200 times. A GC value with a channel pair above the 95^th^ percentile of the null distribution was considered as significant. The spectral GC values from both directions across 1-100 Hz were then averaged across all pairs for each participant.

### Phase-amplitude coupling

We analyzed the PAC on channel pairs between the hippocampus and the EC in each participant. To extract the directionality information from each pair, the relationship between a lower-frequency (2-30 Hz) phase (in steps of 1 Hz) from the hippocampus/EC channel and higher-frequency (30-100 Hz) amplitude (in steps of 2 Hz) from the EC/hippocampus channel was examined individually for correct and incorrect trials by calculating the mean vector length (57). Then, the raw PAC was assessed against surrogate distributions via bootstrapping. Specifically, we shuffled the data by inserting a randomly generated time lag between the time series of the low frequency phase and the high frequency amplitude and recomputed the PAC. This procedure was repeated 200 times to create surrogate distributions of PAC values. Then the raw PAC was *z*-scored based on the null distributions to correct for any spurious results. For each participant, the *z*-scored PACs from the two directions were averaged across all pairs between the hippocampus and the EC.

### Spectral power

Time-frequency power was separately computed for correct and incorrect trials. For each trial and each channel, we convolved the signal with complex-valued Morlet wavelets (6 cycles) to obtain power information at each frequency from 1 to 100 Hz (1-30 Hz in steps of 1 Hz and 30-100 Hz in 2 Hz steps) with a time resolution of 1 ms. The task-induced power was analyzed per trial using a statistical bootstrapping procedure (methods have been described in more detail in a previous paper (5)). Then, the raw power for each time point during the task was *z*-scored by comparing it to the null distribution to generate the *z*-scored power. For each participant, the *z*-scored spectral power was averaged across the maintenance period within the hippocampus and the EC separately for the correct and incorrect trials.

### Statistical analysis

We used a cluster-based permutation test (58) to examine whether the spectral GC values differed between the two directions (from the hippocampus to the EC and the reverse direction) at frequency points at the group level for correct and incorrect trials, separately. We created a Monte Carlo distribution by randomly shuffling the labels of the directions 1000 times. The differences between the directions were considered significant when the maximum of the cluster-level summed *t* values in the true data exceeded the 95^th^ percentile of the null distribution (*p* < .05).

Next, we extracted the theta phase (2-8 Hz) - gamma amplitude (30-100 Hz) *z*-scored PAC from two directions: the theta phase of the hippocampus modulating the gamma amplitude of the EC as well as the theta phase of the EC modulating the gamma amplitude of the hippocampus for the correct and incorrect trials, separately. To assess the real coupling effects, the *z*-scored PACs were compared with 0 using a one-tailed cluster-based permutation test used in the GC analysis. Higher values relative to 0 were defined as real coupling (*p* < .05). In addition, given the *z*-scored power analysis, the *z*-scored powers at the theta and gamma bands were separately extracted from the hippocampus and the EC for the correct and incorrect trials. Next, we made comparisons of the *z*-scored power between the correct and incorrect trials using paired *t* tests separately for the theta and gamma bands for the two regions. We considered *p* < .05 as significant.

### Machine learning analysis

We used multi-variate pattern analysis (MVPA) to decode the WM outcome (correct or incorrect) from the neural activity within the hippocampal-EC circuit during WM maintenance. Following our hypothesis, we used the patterns from the GC, *z*-scored PAC, and *z*-scored power within the hippocampus and EC as our features. In this study, we used SVM (59) as a classifier to classify the WM outcome. SVM is widely used in decoding analyses in neuroimaging studies (8) because of its suitability for analyses with a relatively small number of samples. It is provided by the COSMOMVPA package (60) in MATLAB. The details of our MVPA decoding analyses were as follows:

(A) GC patterns: We considered the GC patterns from two directions, from the hippocampus to the EC and the reverse direction, to allow us to investigate whether there was a specific information flow pattern that could decode the WM performance. The GC patterns were extracted at each frequency band: theta (2-8 Hz), alpha (8-13 Hz), beta (13-30 Hz), and gamma (30-100Hz) (**Fig. 2(B)**) from trials with correct or incorrect performance, separately. For each participant and each outcome, the GC pattern included *M* values (*M* = 25 in theta, *M* = 21 in alpha, *M* = 69 in beta, and *M* = 281 in gamma) and these values were converted into a feature vector. We trained an SVM classifier with a linear kernel with the cost equal to one. To increase the impact of the analysis to a larger population, we merged the data from all the participants (*N* = 13) and performed the classification across the participants. Specifically, we used the feature vectors labeled as correct and incorrect from *N*-1 participants as a training dataset (*N*-1 * M features * 2 outcomes) and tested these on the remaining dataset. We did an inter-participant cross validation by leaving one participant out for validation and replicated the classification process *N* times. The schematic of the MVPA using the feature patterns is shown in **Fig. 1(C)**. The accuracy of the classifier was averaged across *N* cross-validations as a measure of performance. This process was repeated for each direction and each frequency band, so we performed 8 (2 directions * 4 frequency bands) classifications.

(B) *Z*-scored PAC patterns: We extracted the theta phase - gamma band *z*-scored PAC for the correct and incorrect trials separately. For each participant and each outcome, there were 7 phase bins * 36 amplitude bins = 252 features converted to a feature vector. We used principal component analysis (PCA) to reduce the number of the features to several principal components (*M*) that explained 95% of the variance in the data. Then, we combined all the participants (*N*)’ data from trials with correct or incorrect performance, fed them into a linear SVM classifier (default parameters), trained the classifier on the dataset from *N*-1 participants (N-1 * M principal components * 2 outcomes), and tested it on the remaining one participant. Similar to the cross-validation performed in the previous analysis, we left one participant out for validation and replicated this procedure *N* times. The accuracy of the classifier was averaged across all replications. In total, we separately performed this classification process on 2 directions: from the hippocampus modulating the EC and from the EC modulating the hippocampus.

(C) *Z*-scored power pattern: We used a frequency specific *z*-scored power pattern at the theta (2-8 Hz, 7 frequency bins) and gamma (30-100 Hz, 36 frequency bins) bands from the hippocampus and EC to decode the WM outcome. The averaged *z*-scored power values at each band within each region were computed, for the correct and incorrect trials, separately. We therefore performed 4 (2 regions * 2 frequency bands) classifications in this decoding analysis. The training dataset for the linear SVM classifier included the data from *N*-1 participants (*N*-1 * M features * 2 outcomes) and the classifier was tested on the remaining one participant’s dataset. The accuracy of the classifier was averaged across *N* replications by leaving one participant out for the cross-validation.

For all the MVPA decoding analyses, we used a nonparametric permutation approach to test the significance of the accuracy values. We created 1000 unique permutations of the true labels of the classifier. A null distribution of the accuracy was generated using the training data with randomized condition labels. The null distribution was generated for each direction, each frequency band, and each region separately. To correct for multiple comparisons across all directions, frequency bands, and regions (total 8 classifications for GC, 2 for *z*-scored PAC and 4 for *z*-scored power), we took the maximum value of the null distribution across all tests, which provided a final null distribution. The original accuracy value (found from a classifier with true labels) exceeded the 99^th^ percentile of the null distribution (*p* < .01) and was considered significant.

## Data availability statement

The data set was analyzed and described earlier (4, 5, 11, 38) and is freely available for download at https://doi.gin.g-node.org/10.12751/g-node.d76994/. The task is freely available for download at http://www.neurobs.com/ex_files/expt_view?id=266. Links to updates and further data sets can be found at https://hfozuri.ch.

## Acknowledgments

The authors appreciate the suggestions of Prof. Congying Chu from the Brainnetome Center in the Institute of Automation, Chinese Academy of Sciences, and Junjie Zhuo from Hainan University. The authors also thank Rhoda E. Perozzi and Edmund F. Perozzi, PhDs, for English and content editing assistance.

